# Systems analysis of immune responses to attenuated *P. falciparum* malaria sporozoite vaccination reveals excessive inflammatory signatures correlating with impaired immunity

**DOI:** 10.1101/2021.07.28.454252

**Authors:** Ying Du, Nina Hertoghs, Fergal J. Duffy, Jason Carnes, Suzanne M. McDermott, Maxwell L. Neal, Katharine V. Schwedhelm, M. Juliana McElrath, Stephen C. De Rosa, John D. Aitchison, Kenneth D. Stuart

## Abstract

Immunization with radiation-attenuated sporozoites (RAS) can confer sterilizing protection against malaria, although the mechanisms behind this protection are incompletely understood. We performed a systems biology analysis of samples from the Immunization by <u>M</u>osquito with <u>R</u>adiation <u>A</u>ttenuated <u>S</u>porozoites IMRAS) trial, which comprised *P. falciparum* RAS-immunized (*Pf*RAS), malaria-naive participants whose protection from malaria infection was subsequently assessed by controlled human malaria infection (CHMI). Blood samples collected after initial *Pf*RAS immunization were analyzed to compare immune responses between protected and non-protected volunteers leveraging integrative analysis of whole blood RNA-seq, high parameter flow cytometry, and single cell CITEseq of PBMCs. This analysis revealed differences in early innate immune responses indicating divergent paths associated with protection. In particular, elevated levels of inflammatory responses early after the initial immunization were detrimental for the development of protective adaptive immunity. Specifically, non-classical monocytes and early type I interferon responses induced within 1 day of *Pf*RAS vaccination correlated with impaired immunity. Non-protected individuals also showed an increase in Th2 polarized T cell responses whereas we observed a trend towards increased Th1 and T-bet+ CD8 T cell responses in protected individuals. Temporal differences in genes associated with natural killer cells suggest an important role in immune regulation by these cells. These findings give insight into the immune responses that confer protection against malaria and may guide further malaria vaccine development.

## Introduction

Malaria is a devastating disease that results in over 200 million cases and hundreds of thousands of deaths annually. *Plasmodium falciparum* causes the most serious disease and the most deaths, especially in sub-Saharan Africa and primarily in children (Vekemans et al., 2021). Multi-pronged efforts to eliminate malaria have led to substantial reductions in malaria incidence but the development of drug and insecticide resistance as well as other factors, including the current COVID-19 pandemic, are a challenge to further progress (Vekemans et al., 2021; Weiss et al., 2021). An effective anti-malarial vaccine has been a long term goal which has proven challenging. Currently, a single approved malaria vaccine exists, the RTS,S subunit vaccine, which elicited 28-33% protection in infants over a 4-year study period (RTSS Clinical Trials Partnership, 2015). An improved vaccine, especially one that prevents infection, would be a valuable tool in the effort to eliminate this disease. Understanding the immune responses that contribute to vaccine induced immune protection could aid the development of such vaccines.

Sporozoites (SPZs) are the liver-infectious life cycle stage of malaria, injected via mosquito bite in natural infections. Many studies in humans and model systems have shown that vaccination with *P. falciparum* SPZs that have been attenuated by radiation, genetic modification, or drug treatment can result in sterilizing immunity, as determined by subsequent controlled human malaria infection (CHMI) (Coelho et al., 2017; Obeng-Adjei et al., 2015; Portugal et al., 2015, 2014). This mode of vaccination aims to elicit immunity against pre-erythrocytic parasite stages, where the biomass of the parasites is low and the infection is asymptomatic. Currently, no universal correlates of protection have been identified and the nature of protective immunity is incompletely understood. Sterilizing immunity is likely to be complex and directed at multiple antigens given that the *P. falciparum* genome encodes more than 5,300 unique proteins. Available evidence indicates that antibodies against major surface proteins of infecting SPZs, e.g. CSP and TRAP, contribute to protection (Ishizuka et al., 2016; Seder et al., 2013). Animal models have indicated that liver-resident CD8 T cell responses are important for protection, which is inherently challenging to study in humans, as the human liver is not readily accessible for sampling and a very small fraction of its cells get infected (Fernandez-Ruiz et al., 2016b; Trieu et al., 2011). It is imperative to identify correlates of protection in humans that can aid the improvement of the current vaccines and development of vaccine candidates. To this end, human vaccination and challenge trials with attenuated *Pf*SPZs provide an opportunity to elucidate immune responses that are associated with pre-erythrocytic protection.

In this study, we applied a systems immunology approach to identify correlates of protection that are identifiable up to 28 days after initial vaccination in malaria naïve human trial subjects that participated in the Immunization by <u>M</u>osquito with <u>R</u>adiation <u>A</u>ttenuated <u>S</u>porozoites (IMRAS) trial (Hickey et al., 2020). Participants were immunized by *Pf*RAS delivered by mosquito bite with efficacy assessed by CHMI. Five total immunizations were delivered, spaced 4 to 5 weeks apart. The trial was designed with a suboptimal vaccine dose regime to elicit approximately 50% vaccine efficacy to facilitate comparison between protected (P) and non-protected (NP) subjects. Of particular interest in the IMRAS trial is the prime vaccination. IMRAS participants are malaria unexposed, and the initial *Pf*RAS vaccination represents the first time their immune system has been exposed to *P. falciparum* sporozoites. We hypothesized that the earliest immune resposnes to *Pf*RAS represent a critical time period determining subsequent development of sterilizing immunity.

Our integrative analysis of whole blood transcriptomics, high parameter flow cytometry and single cell CITE-seq identified numerous vaccine-induced responses including ones that correlated with protection. We observed strong negative correlations with protection that were associated with inflammation, type I interferon (IFN), and signatures related to monocytes and neutrophils, and type 2 polarized T helper cell responses. Differential kinetics in natural killer (NK) cell-associated responses and a trend of increased T-helper 1 cells correlated positively with protection. These results suggest that the priming vaccination with radiation attenuated *Pf*SPZs establishes immunological trajectories that can result in protection following additional vaccinations, and show that early inflammatory responses can negatively influence the fate of protective immunity.

## Results

The IMRAS cohort analyzed consisted of eleven malaria-naïve adults immunized with five doses of approximately 200 bites from *Pf*RAS NF54 infected mosquitos (Hickey et al., 2020). The first four doses were given four weeks apart and the final dose was administered five weeks after the fourth. Protection was tested by controlled human malaria infection (CHMI) three weeks after the final vaccination (Fig. S1 A). Six of the eleven immunized participants were protected, i.e. zero parasitemia after CHMI. Of the five non-protected subjects one developed parasitemia on day 9 after CHMI and four did so on day 13 after CHMI (Fig. S1 B). There was no significant correlation between the number of *Pf*RAS infectious mosquito bites received and protection status among true-immunized subjects (Fig. S1 C).

### PfSPZ vaccination induced broad transcriptome responses

Whole blood transcriptome profiling was performed on all eleven immunized IMRAS participants at 6 timepoints after the initial *Pf*RAS vaccination. We examined transcriptional changes between adjacent timepoints, which we refer to as time “intervals”, namely between days 0-1, 1-3, 3-7, 7-14, and 14-28 after immunization. We conducted linear mixed-effects regression modeling analysis (LMER) of the responses to identify significantly responsive genes over all subjects and those that differed between P and NP at each interval (FDR < 0.2 & p < 0.05; see Materials and Methods). 90% confidence intervals (CIs) were calculated around model coefficients to label genes as either increased or decreased if the CI was entirely above or below 0, respectively (Fig. 1A). Many significantly responsive genes were observed after the first *Pf*SPZ vaccination; 8170 genes had increased or decreased expression responses over at least at one interval accross all immunized subjects and approximately 10% of these differed significantly between P and NP subjects (Fig. 1A).

**Figure 1.**
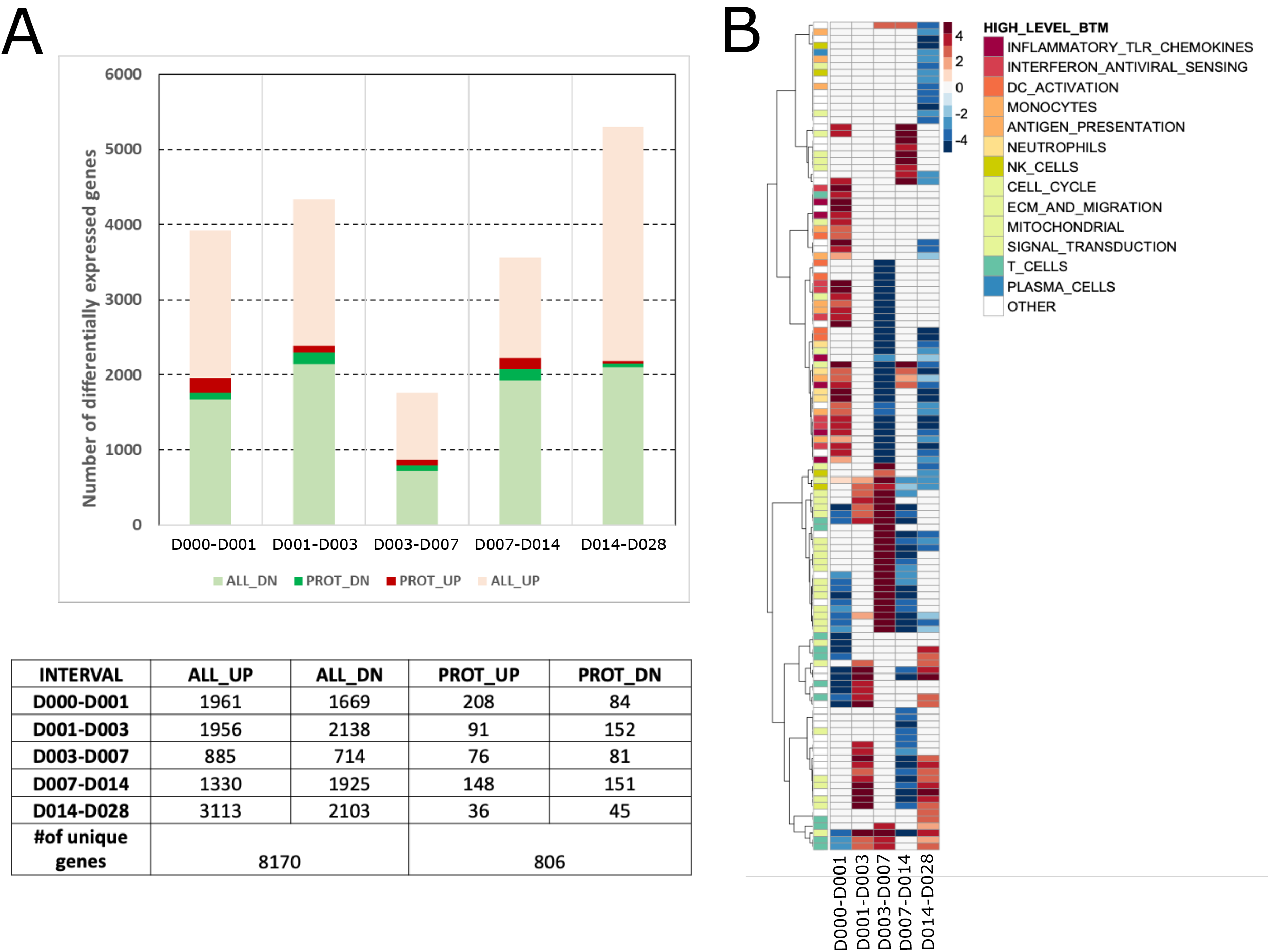
Vaccine induced gene responses after the first immunization. **A** Barplot and table showing numbers of vaccine-induced genes with increased (red) or decreased (green) expression over each time interval in all immunized subjects (ALL_UP, ALL_DN) (FDR < 0.2, p <0.05, 90% CI > 0 or < 0). Darker colors indicate genes that differ significantly in expression between protected (P) and non-protected (NP) subjects (PROT_UP, PROT_DN). **B** Heatmap of modules significantly enriched for vaccine-induced genes. Each row represents a BTM, each column represents a time interval. Heatmap color shows hypergeometric effect size (ES) of a BTM enriched in genes with increased (red/positive ES) or decreased (blue/negative ES) expression. Non-significant BTMs are shown in white. Assignment of a BTM to a high-level annotation group is illustrated by a colored sidebar.

The association between vaccine induced gene responses and specific cell populations and immunological processes was determined by testing whether pre-defined coherent blood transcription module sets (BTMs) (Chaussabel et al., 2008a; Li et al., 2014a; Liberzon et al., 2015) showed enrichment for significantly responsive genes (hypergeometric p < 0.1). We identified 122 BTMs significantly enriched in response genes in at least one time interval. Hierarchical clustering of enriched BTM hypergeometric effect sizes at each interval revealed discrete time-dependent response patterns among the immunized subjects (Fig. 1 B). BTMs increased during the first day after vaccination were associated broadly with immunity and inflammation, including TLR sensing; antigen processing; interferon and inflammation; monocytes and neutrophils. This was accompanied by decreases in BTMs associated with the cell cycle and T cells. Relatively few enriched BTMs were observed between day 1 and 3. Over subsequent intervals (D3-7, D7-14, D14-28) BTMs associated with monocytes, neutrophils, TLR sensing, inflammation and interferon, decreased sharply. However, we observed an increase in cell cycle-associated responses from D3-7, and an induction of T cell associated BTMs at D14-28. Overall, the priming vaccination resulted in robust and dynamic transcriptional responses in the combined group of P and NP subjects.

#### Protection associated genes showed distinct response dynamics in P and NP individuals

To further explore reponse dynamics of protection-associated genes, we performed unsupervised clustering of the responses over time of the 1394 genes that were differentially expressed between P and NP (FDR < 1/3 & p < 0.05, Fig.2 A; Materials and Methods). Hierarchical clustering was performed separately for P and NP subjects to illustrate the response differences between these two groups and to identify sets of genes with coherent response profiles over time following *Pf*SPZ immunization (Fig. 2 A,C). Immune functions and molecular mechanisms associated with these clusters were identified by the hypergeometric enrichment test using BTMs and by Ingenuity Pathway Analysis (IPA) (see Materials and Methods) (Fig. 2 B,D, Fig. S4). We identified four major gene clusters for P subjects (P_1, P_2, P_3, P_4) and five for NP subjects (NP_1, NP_2, NP_3, NP_4, NP_5). The patterns in which these 1394 genes changed over time differed between P and NP, leading to substantial differences in gene composition of most of the major P and NP clusters (Fig.2 A,C,E). A total of 39 signficantly enriched BTMs and 159 significant IPA pathways were identified between P and NP (Table S1).

**Figure 2.**
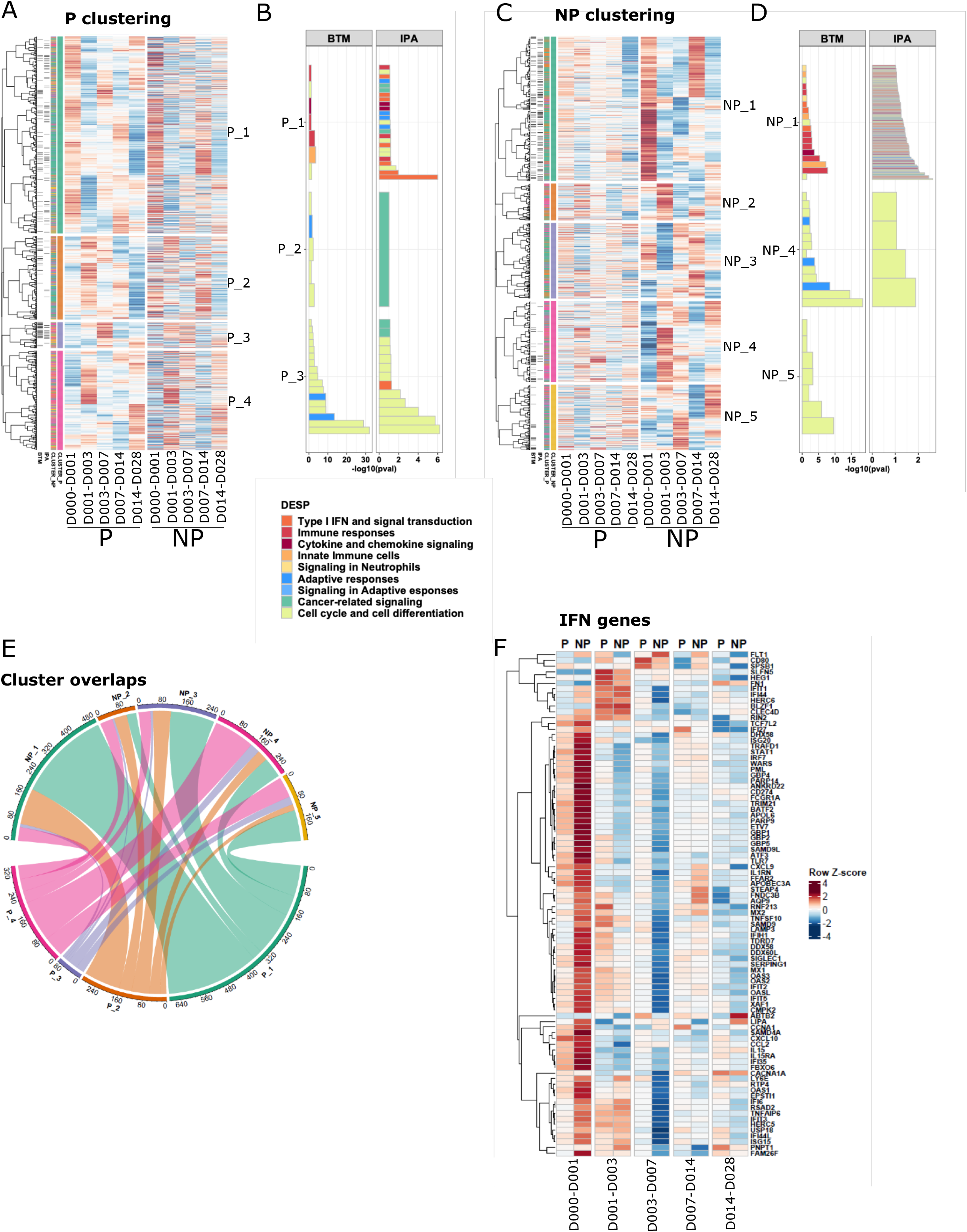
Vaccine induced protection associated gene responses. **A**,**C**. Heatmaps showing 1394 protection-associated genes ordered by hierarchical clustering based on expression in P (**A)** and NP (**C**) subjects for each time interval. Black sidebars indicate genes associated with an IPA pathway or BTM. Colored sidebars indicate P and NP gene clusters. Expression values were z-score transformed in rows for visualization. **B**,**D**. Enriched BTMs and IPA pathways for clusters from P (**B**) and NP (**D**) subjects. X axis represents –log10(FDR) generated from hypergeometric tests. Color indicates assignment of a BTM module or IPA pathway to a high-level annotation group. **E**. Circos plot showing overlap between P and NP clusters, numbered to match **A** and **C**, and colored by cluster number (e.g: P_1, NP_1: green, P_2, NP_2:orange). **F**. Expression changes of type I interferon-associated genes in P and NP subjects over each interval.

Most gene clusters from both P and NP were strongly enriched for cell cycle-associated BTMs (Fig.2 B,D). An important exception was P_1 and NP_1, both were associated primarily with various immune response modules including many type I IFN-associated modules. These two gene clusters have the largest total numbers of genes and associated BTMs. In total, P_1 and NP_1 have 317 genes in common (hypergeometric p = 4.2e-17) (Fig. 2 E), although numbers of enriched modules and gene expression dynamics are distinct (Fig. S2 A-C). Expression of most genes in these clusters was increased over the interval D0-1, with higher response magnitude in NP_1 (Fig. S2 C). Consistent with strong enrichment of type I IFN associated modules in P and NP-associated cluster 1, we observed that IFN-stimulated genes (ISGs) were strongly increased overall in NP compared with P and NP by day 1 after the first vaccination (Fig.2 F, Mann Whitney U test, P < NP, p < 2.2e-16) (Kazmin et al., 2017). Furthermore, we observed significantly increased expression of genes related to sensing through Pattern Recognition Receptors (PRR) in NP participants by day 1, for both MyD88 dependent and independent pathways (Fig. S3 A-C). Altogether, these patterns indicate that overall innate sensing and inflammatory responses were highly elevated in NP compared to P subjects, strongly suggesting that they negatively affected the induction of adaptive immunity against SPZ challenge.

#### Gene set enrichment analysis identified early inflammation as correlate of impaired immunity

To more broadly explore immunological processes and cell types associated with protection we performed gene set enrichment analysis (GSEA) separately for P and NP subjects at each time interval (Fig. 3 A). GSEA facilitated identifying important coherent response modules that our above analysis based on differentially expressed genes may not have revealed. GSEA revealed 80 BTMs significantly enriched at one or more timepoints (FDR < 0.1). Twenty of these BTMs overlapped with the 39 BTMs enriched in differentially expressed genes.

**Figure 3.**
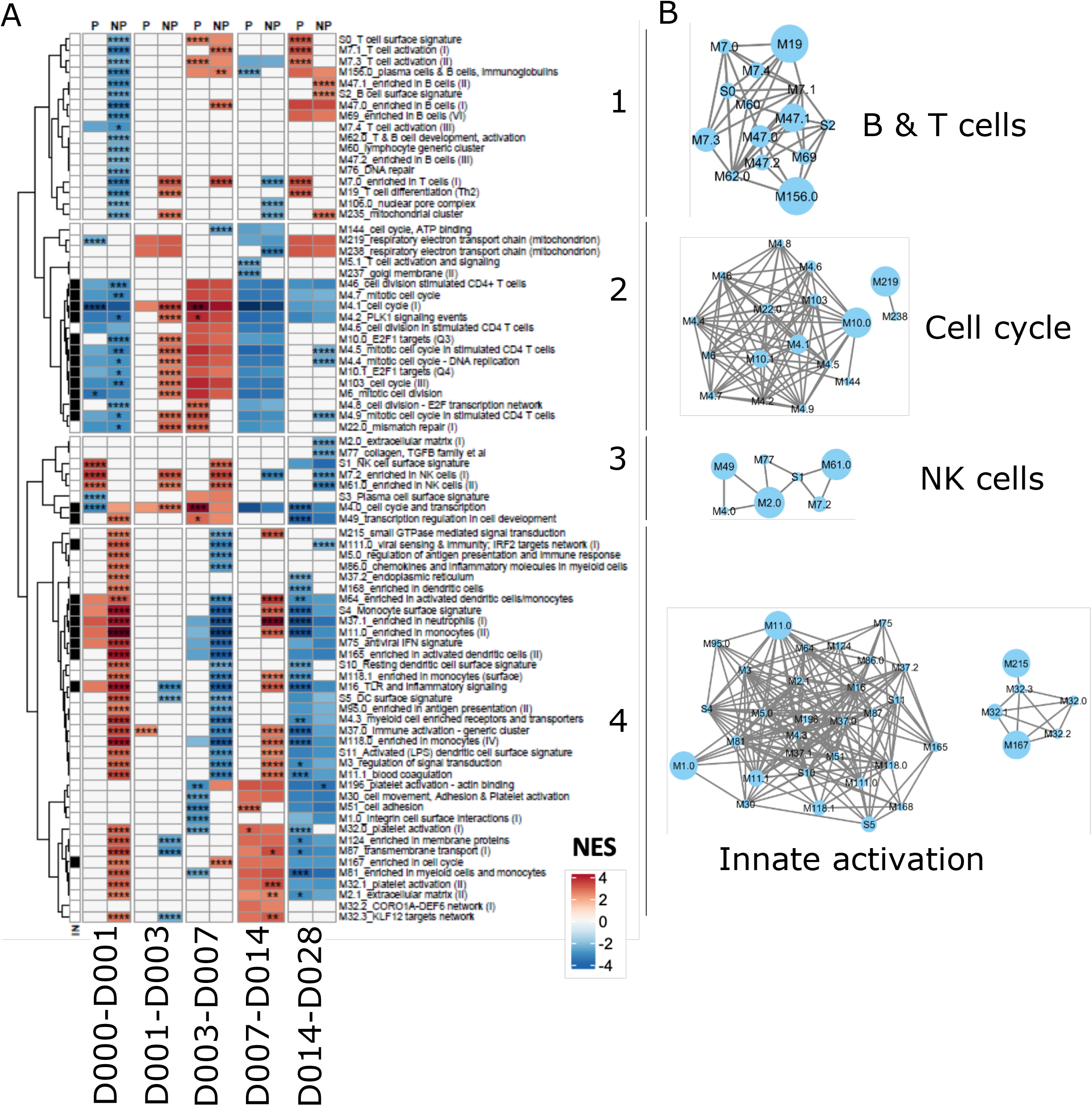
Gene set enrichment analysis after the first immunization in P and NP subjects. **A**. Heatmap of GSEA normalized enrichments scores (NES) derived from BTM expression changes over each time interval. Red represents activated BTMs and blue represents down-regulated BTMs. Asterisks represent significant differences in expression between P and NP for a time interval (Mann Whitney U test. ^****^: p < 0.0001; ^***^: p < 0.001; ^**^: p < 0.01; ^*^: p < 0.05). BTM clusters derived from hierarchical clustering are indicated by numbers to the right of the heatmap. Black sidebar on the left represents BTMs also enriched in unsupervised clusters of protection associated genes in P and NP subjects (Fig 1 B). **B**. Relationships between BTMs within the four identified clusters. BTMs with shared genes are connected by lines and the node sizes correspond to the numbers of shared genes. Predominant cell-type module annotations for each cluster are shown.

Patterns of responses over time differed between P and NP. Two modules showed opposite responses: *M4*.*0 cell cycle transcription* over day 0-1, and *M196 platelet activation + actin binding* over day 3-7 were both increased in P and decreased in NP. Hierarchical clustering of the GSEA BTM normalized enrichment scores (NESs) identified four major clusters of BTMs (Fig. 3 A) and many of the BTMs within a cluster shared genes (Fig. 3 B). GSEA cluster 1 is associated with B and T cells, cluster 2 with cell cycle and potentially cell proliferation, cluster 3 with NK cells, and cluster 4 with monocytes, neutrophils and immune activation.

Cluster 1 contained BTMs associated with T and B cells. These BTMs decreased over day 0-1 specifically in NP, with most BTMs remaining unchanged at all other time intervals for both P and NP. BTMs in cluster 2 were generally associated with cell cycle and division. These BTMs were downregulated over day 0-1 and day 7-14 and upregulated from day 3-7 in both P and NP.

Notably, most of these BTMs were activated in NP but not in P from day 3-7. Cluster 3 was the most heterogeneous cluster in terms of composition and time dynamics, and included NK cell, plasma cell and trancriptional regulatory BTMs among others. These modules tended to be increased at early time intervals, before decreasing at day 7-14 and day 14-28. Notably, NK BTMs were upregulated in P over day 0-1; however, their activation occurred late in NP over day 1-3 and day 3-7. Cluster 4 primarily represented innate inflammation, interferon, monocytes, neutrophils and dendritic cell BTMs. Consistent with our above gene-based analysis, cluster 4 responses in NP individuals increased sharply at day 0-1 and day 7-14 with an intermediate decrease from day 3-7, with relatively few changes in P individuals over the first four time intervals. Both P and NP individuals showed decreased responses in cluster 4 BTMs day 14-28. Cluster 4 also included platelet BTMs, and early activation of platelets in NP may reflect their influence on the elevated expression associated with immune activation and monocytes (Rolfes et al., 2019). Activated neutrophils produce reactive oxygen species (ROS), and we observed an increase in ROS signalling in NP (Fig. S5 A-C) at day 0-1. The differences between P and NP subjects in this cluster highlights the greater magnitude of inflammatory responses by innate cells in NP versus P subjects early after the first immunization. Overall, this analysis supports our hypothesis that high levels of inflammatory responses and type I IFN are detrimental for protective immunity, and suggests these responses are associated with monocytes, DCs and NK cells.

#### Specific Immune cell types associated with protection by trancriptomics and flow cytometry

GSEA and differential gene expression analyses indicated that specific immune associated transcriptional responses were induced at different times and to different extents between P and NP subjects. This suggests that specific immune cell populations responded differently in P vs NP, thus, we further investigated whether a cell type-specific signature could be identified after initial *Pf*RAS immunization.

We generated heatmaps of expression changes by averaging expression of genes from significant cell-type associated BTMs identified by the GSEA analysis (Fig. 4 A,B, Fig. S6 A-G). Monocytes, DCs, neutrophils and platelet-associated responses were highly increased in NP over day 0-1, indicating strong innate immune responses in NP that were absent or significantly lower in P subjects (Fig. 4 A,B, Fig. S6 A-D). In contrast, genes associated with T and B cells were strongly decreased over day 0-1 in NP subjects (Fig. 4 A, Fig. S6 E-F). Both P and NP subjects had a modest increase in B cell associated genes in the last time interval (Fig. S6 F). Interestingly, P and NP subjects differed in the timing, magnitude and nature of active NK cell-related genes (Fig. 4 A). Once again, these results indicate that early innate immune activation strongly correlates with insufficient immunity to challenge after the completion of the vaccine regimen and transcriptome responses associated with adaptive cell types show the opposite expression pattern.

**Figure 4.**
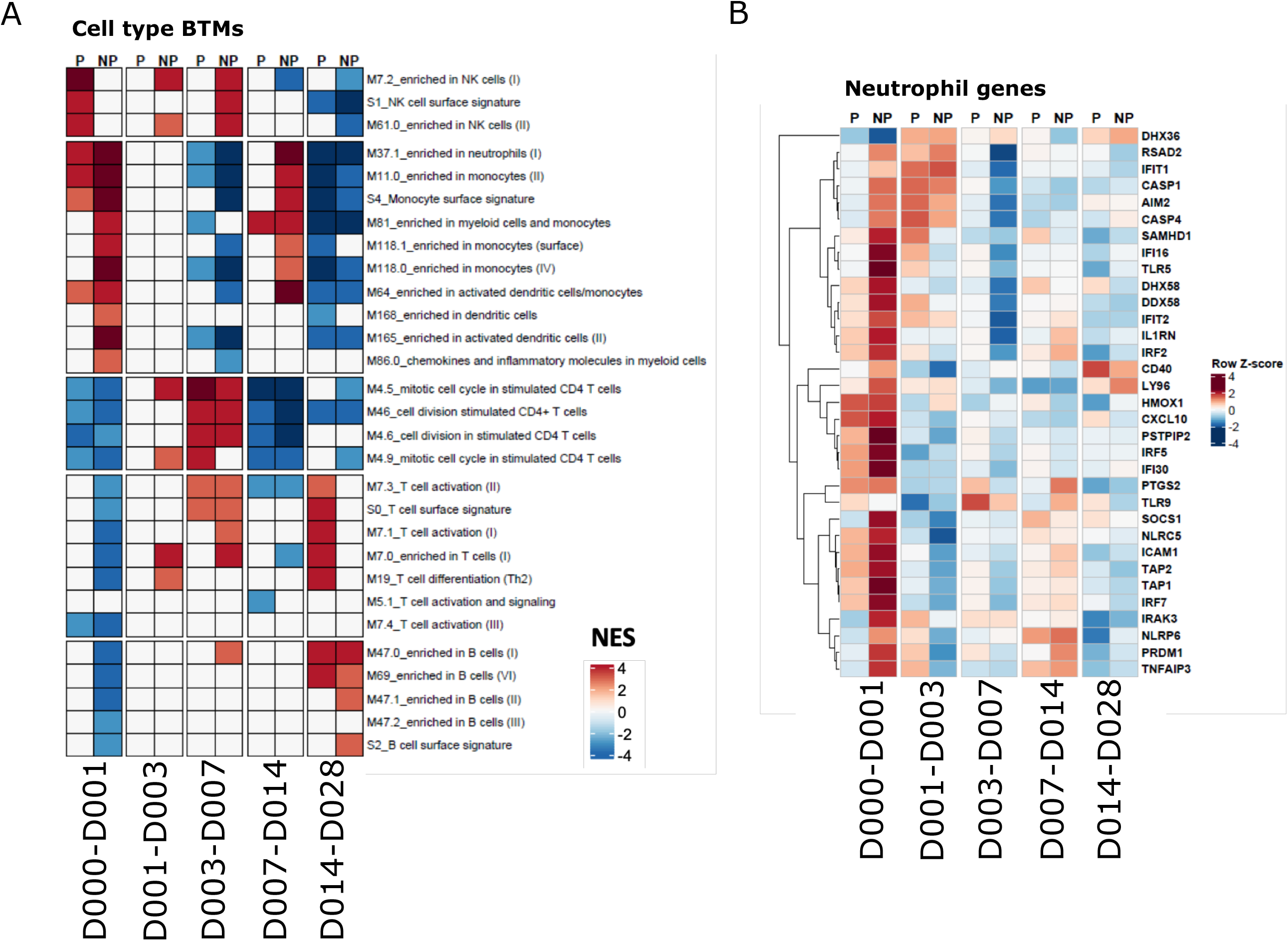
Cell type-associated responses after the first vaccination. **A**. Heatmap showing cell type specific BTM GSEA NESs in P and NP subjects. **B**. Heatmap showing expression changes of malaria responding neutrophil genes in P and NP subjects on the sampling day compared with the previous sampling day. Expression values were z-score transformed by row for visualization.

To validate this whole blood transcriptional analysis, we applied high parameter flow cytometry to characterize PBMCs isolated from immunized individuals after the first immunization. This was done at overlapping time points with the whole blood transcriptional analyses, although we lacked matching PBMC samples for the 1 day and 28 days post immunization timepoints (Table S2, Fig.5 A-G). We assessed whether transcriptional responses associated with specific cell types were correlated with flow cytometry derived counts of the appropriate populations.

**Figure 5.**
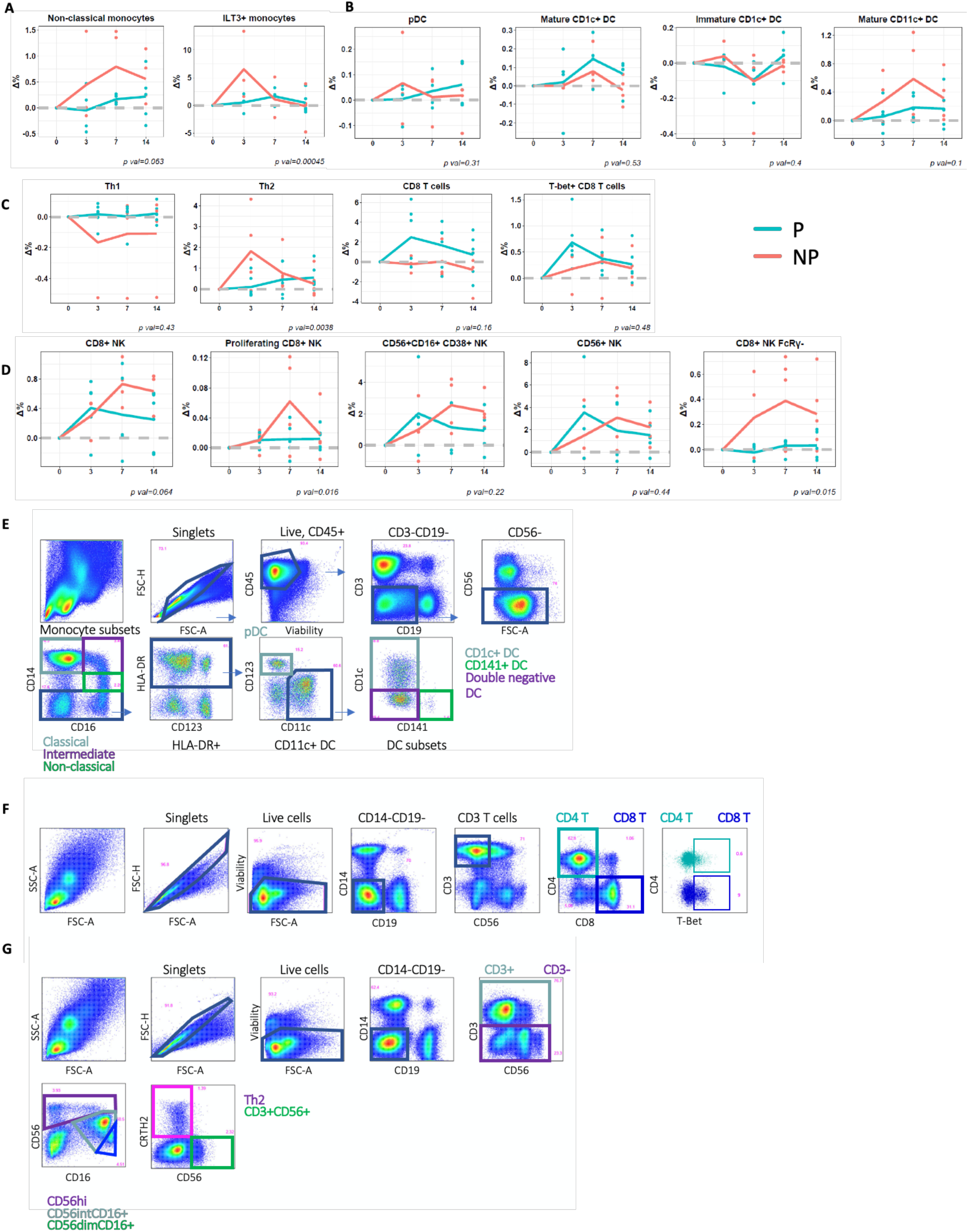
Temporal changes of specific cell types across time intervals in P and NP subjects. **A-D**. Changes in flow-cytometry measured PBMC sub-populations over time, relative to day 0. **A**. non-classical monocytes and ILT3+ monocytes **B**. DCs **C**. T cells **D**. NK cells. Points represent cell proportion values in each subject. Line represents average values across P and NP subjects. LMER p-values are shown for P vs NP differences. **E-G**. Flow cytometry gating schemes **E**. DC subsets and monocytes **F**. Th1 and Th2 cells **G**. NK cells.

Consistent with the RNAseq data, NP subjects showed a trend towards increased numbers of circulating monocytes shortly after the first immunization, specifically non-classical monocytes (CD14+-CD16+) (Fig5A, p = 0.06). These inflammatory cells have been implicated in several vaccination studies to be correlated with impaired immunity (George et al., 2018; Mitchell et al., 2012). We also observed increased numbers of ILT3+ monocytes, associated with IFN exposure (Jensen et al., 2010; Waschbisch et al., 2014), after the first vaccination (Fig.5 A, p = 0.0004). Furthermore, by applying a previously identified transcriptional signature of macrophage polarization (Becker et al., 2015), we observed that responses associated with classically activated macrophages were specifically increased in NP between day 0-1, but no change was observed for alternatively activated macrophage responses (Fig S6 H,I). We compared the abundance and activation status of several DC subsets between P and NP subjects and found several trends that were consistent with RNA-seq results, albeit not statistically significant (Fig. 5 E). We observed a subtle increase in plasmacytoid DCs, and overall increased CD11c+ DC numbers 3 days after the first immunization in NP compared to P subjects (Fig. 5 B). Interestingly, the proportion of “mature” CD1c+ DCs expressing CD86 was higher in P than NP subjects whereas CD86 expressing CD11c+ DCs that lack CD1c or CD141 expression, and non-activated CD1c DCs were enriched in NP subjects. These findings suggest that CD11c+ DCs, non-activated CD1c+ DCs and non-classical monocytes contribute to the transcriptional signature in NP subjects where we observed an increase in DC and monocyte-associated genes (Fig.5 B, Fig. 4 A, Fig. S6 B,H,I). Because different types of DCs can activate and skew different T-helper cell responses, we also assessed the proportion of Th1 and Th2 CD4+ T cells (Collin and Bigley, 2018) and observed greater proportions of Th2 CD4 T cells in NP, and a trend towards more Th1 cells in P subjects (Fig. 5C). In addition, we found a trend of higher circulating numbers of CD8 T cells and T-bet expressing CD8 T cells in P subjects. In CD8 T cells, T-bet is an important transcription factor that is involved in memory cell formation (Sullivan et al., 2003). These data suggest that differences in innate responses contributed to impaired protective immunity by skewing the CD4 T cells toward a type 2 phenotype, and protective responses are hallmarked by Th1 and CD8 T cell responses. The abundance of the NK cells in general, and CD38 expressing NK cells matched the transcriptome responses in which NK-associated gene increases peaked at day 1-3 in P subjects and day 7 in NP subjects (Fig. 5D). However CD8-expressing NK cells subsets did not show any early peak in P participants. Notably, CD8+ NK cells lacking FcRγ were more abundant in NP subjects across all measured timepoints (Hart et al., 2019; Hwang et al., 2012; McKinney et al., 2021; Zambello et al., 2020).

To quantify the relationships between the transcriptional and cytometric data, we performed correlation analysis of temporal changes of BTMs and cell subsets from manually gated flow cytometry data (Fig. 6 A). These analyses showed that transcriptional activation of the innate immune response (inflammatory responses, monocyte signatures, DC signatures, viral sensing & immunity, antigen processing & presentation, etc.) positively correlated with increases in the proportion of innate immune cells (DCs, monocytes) (Fig. 6 B). In contrast, transcriptional down-regulation of T cell-related responses (T cell activation, T cell differentiation) paralleled cellular decreases of CD3 T cells (Fig.6B), and transcriptional activation of several NK-associated genes positively correlated with increases in NK cells (Fig. 6 B). The activation of innate immune responses (increases in BTMs related to monocytes, DCs, inflammatory responses) in NP subjects correlated with decreases in total T cell counts (Fig. 6 C). The down-regulation of T cell modules (T cell activation, T cell differentiation) was associated with increases in the abundance of DCs and monocytes (Fig. 6 C). Additionally, the downregulation of B cell-associated genes was correlated with cell composition increases in NKT-like cells that co-express CD3 and CD56 (Fig. 6 C). These analyses indicate that the transcriptional data and the flow cytometry data align consistently and reveal cell abundance and activation changes that correlate with each other.

**Figure 6.**
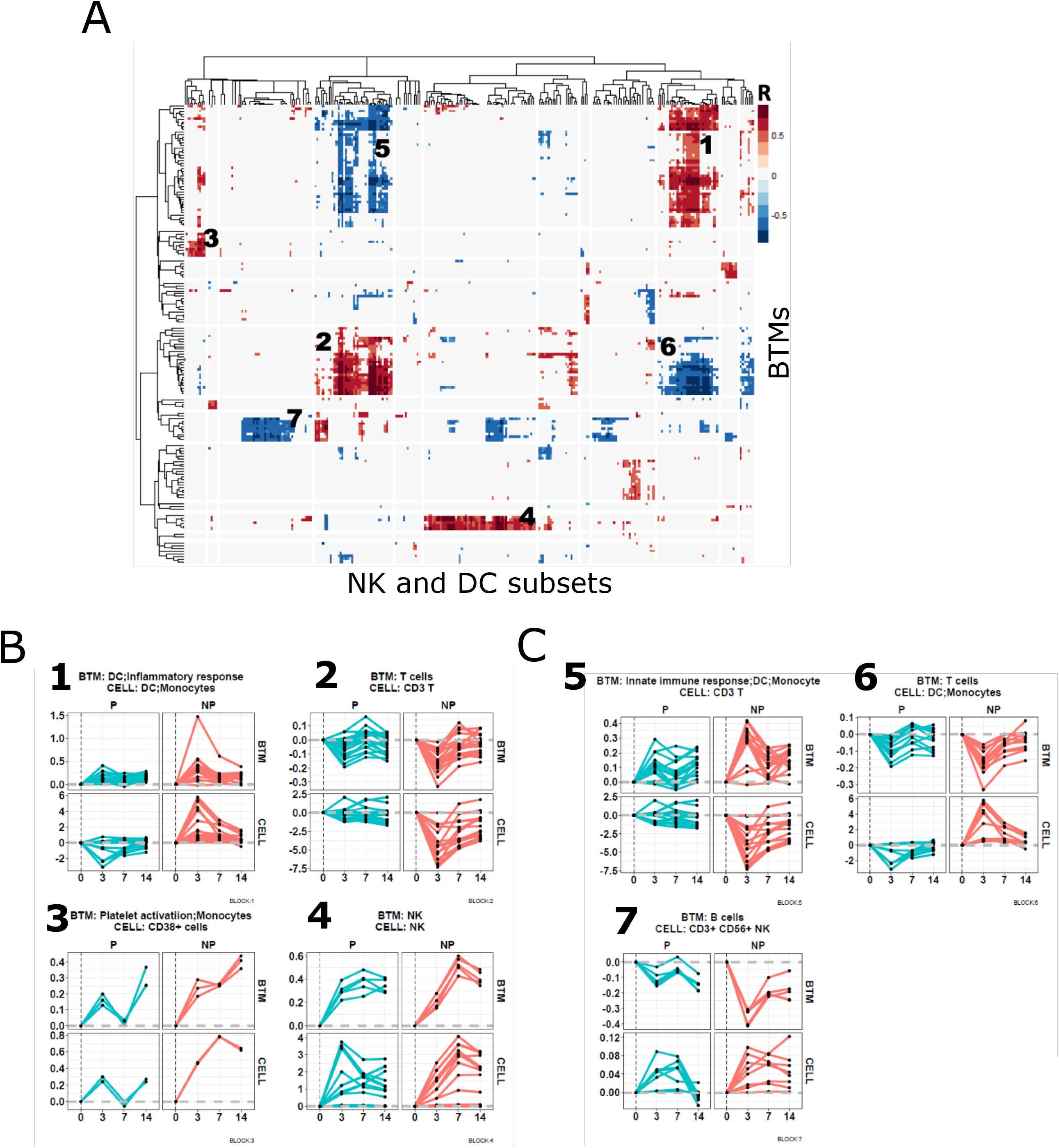
Correlation of temporal changes between cell subsets and BTMs. **A**. Heatmap showing correlations between flow-cytometry derived cell subset counts from NK and DC panels (columns) and BTM expression (rows) matched by participant and time-point. Red represents positive correlation, blue represents negative correlation, and white represents non-significant correlations. Regions of positive and negative correlation are numbered 1-7. **B**,**C**. Changes over time (relative to day 0) of BTMs and cell subsets that are strongly positively correlated (**B**, blocks 1-4 in heatmap **A**) and negatively correlated (**C**, blocks 5-7 in heatmap A). Each line represents average expression changes (relative to day 0) of a cell subset or BTM in P and NP subjects.

#### RNAseq and flow cytometry findings were further validated with scRNAseq

To further validate the identity of cell subsets indicated by transcriptional analysis, we performed CITEseq with a panel of 14 antibodies using PBMC samples collected on day 0, day 3 and day 14 after the priming dose of *Pf*SPZ from an IMRAS subject who dropped out of the trial (Table S3). We obtained single cell RNA seq gene expression and surface protein marker profile data from a total of 12,442 cells. Mapping of these cells to a previously defined multimodal cell atlas based on reference clusters identified 29 cell types in the merged samples (Fig. 7 A) (Hao et al., 2021). We then derived transcriptional cell-type associated signatures using genes highly expressed in CITEseq-identified cell types in our samples, and applied these signatures in GSEA analysis of the whole blood RNA-seq data (Fig. 7 B). Consistent with previous BTM-based GSEA analyses, GSEA with our CITEseq signatures indicated that CD14+ and CD16+ monocyte subsets were highly activated on day 1 in NP subjects. By contrast, genes associated with NK cells were highly activated on day 1 in P subjects, whereas the activation continued on day 3 in NP subjects (Fig. 7 B,C). Finally, comprehensive correlation analysis was performed to examine the relationship between cell subsets predicted by CITEseq and cell-specific BTMs identified in our previous GSEA (Fig. 3, Fig. 4). The high correlation between cell-specific BTMs and corresponding cell types identified in CITEseq data demonstrates that the cell subsets identified by both approaches are consistent (Fig. 7 D) and supports our interpretation of these responses. Thus, integration of whole blood RNA-seq, high parameter flow cytometry and scRNA-seq of individuals after initial *Pf*RAS vaccination revealed a consistent picture where inflammatory transcripts and non-classical monocytes, and Th2 signalling are specifically increased in NP individuals in the first day after *Pf*RAS vaccination while P individuals are marked by increased early NK-cell and Th1 signalling.

**Figure 7.**
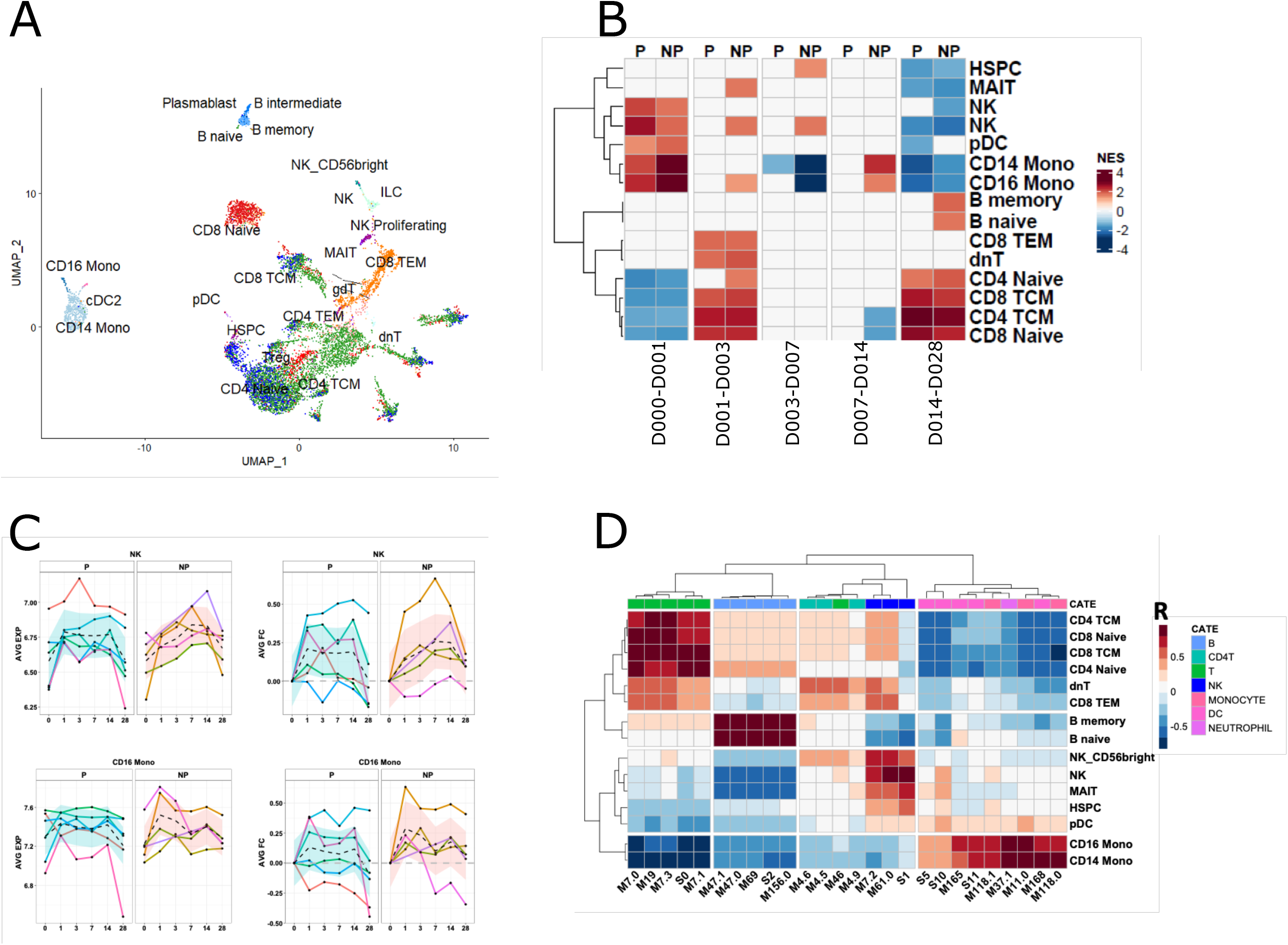
Validation of transcriptional changes of specific cell types using CITEseq data. **A** Scatter plot illustrating cell types identified in CITEseq data, visualized using uniform manifold approximation and projection (UMAP). Each point represents a cell and is colored by cell type. **B** Heatmap showing GSEA NES of CITEseq-derived cell signatures in whole-blood RNAseq in response to the first immunization in P and NP subjects. **C**. Lineplot showing kinetics of CITEseq-identified cell signatures in P and NP subjects. Each line represents the median gene expression levels of a CITEseq gene signature per-subject, and the black dashed line represents the median gene expression levels across all in P or NP subjects. **D**. Heatmap showing Spearman’s rank correlation between cell-type specific BTMs and CITEseq identified cell-specific signatures. The color row bar indicates cell-type annotation of BTMs.

## Discussion

Whole blood transcriptome, high parameter flow cytometry and CITEseq systems analyses of the IMRAS trial identified numerous responses to the first vaccination, including early inflammatory responses that correlated with a lack of protection. These early inflammatory responses were associated with myeloid cells, including neutrophils and increased levels of monocyte and DC subsets, and Th2 cells. This substantial early inflammatory stimulation may have resulted in skewing toward a type 2 response. In contrast, effective protection was correlated with early responses associated with NK cells, later responses associated with T cells and lower overall responses. These results underscore the influence of early innate and adaptive responses on the subsequent immunological trajectories as shown by different response profiles that do or do not ultimately lead to protection from infection (Figs. 2,3). We hypothesize that differential priming of the adaptive compartment by PfRAS vaccinations impacts subsequent responses and ultimately protection from infection by protected vs not protected vaccinees. Thus, immune responses to the first vaccination may be decisive for the outcomes of vaccinations with attenuated sporozoites and may provide early biomarkers of protection.

Myeloid cell activation as a negative correlate of protection may seem counterintuitive since activation is required for antigen presenting cell (APC) priming of adaptive T cell responses in lymphoid tissues. The higher levels of inflammatory responses that we found to correlate with a lack of protection in this study imply that over-induction of inflammation negatively impacted the development of protective adaptive responses. Previously, innate pathways that suppress adaptive responses have been identified in the context of immune pathology (Minkah et al., 2019; Wherry et al., 2007). Similarly, the early increase in neutrophil-associated gene expression following vaccination correlated with a lack of protection. Although neutrophils are rapid innate responders to infection and function in pathogen clearance and immune modulation (Mantovani et al., 2011), a subset of neutrophils that are systemically induced upon acute inflammation suppress T cell responses through ROS production (Kusmartsev et al., 2004; Pillay et al., 2012; Zemans, 2018). The increased expression of transcripts associated with ROS production observed in NP subjects implies that neutrophil suppression of T cell responses may also have negatively impacted development of protective adaptive immune responses after PfRAS vaccination.

Innate priming of naïve T cells into various T-helper cells can occur through the actions of neutrophils and basophils, DC subsets via their intrinsic properties or their interplay with other innate cells, including the induction of Th2 cells in response to type 2 innate lymphoid cell (ILC2)-derived cytokines (Kim and Kim, 2018; Otsuka et al., 2013; Phythian-Adams et al., 2010; Tang et al., 2010; Tjota and Sperling, 2014). The reciprocal early relative increase in Th2 cells and decrease in Th1 cells in NP subjects only implies that inhibition of polarization of Th1 cells by Th2 cytokines suppressed development of responses that protect against CHMI (Kim and Kim, 2018).

We also observed a trend towards higher numbers of Th1 and CD8 T cells in P vs NP subjects at early timepoints, suggesting that Th1 responses contribute to PfRAS vaccination-induced immunity. This is consistent with studies in animal models which concluded that sterilizing protection involves Th1 cells which secrete IFNγ and, in co-ordination with with macrophages and liver resident cytotoxic CD8 T cells, eliminate intracellular pathogens (Cockburn et al., 2013; Fernandez-Ruiz et al., 2016a; Hickey et al., 2020; McNamara et al., 2017; Tse et al., 2014)(Perez-Mazliah and Langhorne, 2014). In addition, the correlation of CSP-specific, IFNγ-producing CD4 T cells with protection from infection following RTS,S/AS02 vaccination (Reece et al., 2004) also suggests that Th1 cells can contribute protective immunity in humans.

Increases in monocyte associated transcriptome responses and of non-classical and ILT3-expressing monocytes in NP subjects suggest that these responses impair development of protection following Pf vaccination. Impairment of vaccine elicited immunity has been linked to non-classical or inflammatory monocytes in mice (George et al., 2018; Li et al., 2013; Mitchell et al., 2012) perhaps involving ILT3 monocyte recruitment to lymph nodes and interference with T cell priming as implied by the enhanced T cell priming following monocyte depletion. GM-CSF production by monocytes and sequestration of cysteine have been suggested as mechanisms of T cell suppression during priming (Mitchell et al., 2012; Serafini et al., 2004). The correlation between type I IFN expression and the lack of protection that observed in this work (Fig 2F) and in a vaccine trial of malaria SPZs given under chemoprophylaxis cover (Tran et al., 2019) indicates that these responses can hamper the development of adaptive responses. That ILT3 surface expression by monocytes can result from type I IFN stimulation (Jensen et al., 2010; Waschbisch et al., 2014) and both type I IFN and ISGs increased expression in NP subjects early after vaccination suggests that the monocytes in NP subjects have encountered type I IFN stimulation. Type I IFNs are potent immune mediators that can directly activate DCs, NK cells and T and B cells and regulate immune responses to many pathogens. These IFNs signal through the interferon alpha receptor (IFNAR) that is present on almost all cells in the body and induce ISGs that function in the control of infection. However, type I IFN has also been linked to immune pathology in chronic viral diseases and in some bacterial infections (McNab et al., 2015). The ultimate protective vs non protective effects of the type I IFN response may well depend on its timing, localization and magnitude of the specific IFN responses as has been suggested (Nagai et al., 2003).

Protection elicited by vaccination with radiation attenuated SPZs requires an abortive liver infection (Doolan and Hoffman, 2000). Infection studies in animals have shown that malaria infected hepatocytes produce type I IFN, and IFNγ-secreting NK and NKT cells are recruited as liver stage parasites are eliminated (Miller et al., 2014). However, type I IFN responses are also detrimental to long term immunity against infection (Liehl et al., 2015, 2014; Miller et al., 2014; Minkah et al., 2019). Caspase-mediated cell death of infected hepatocytes is required for the uptake and presentation of malaria antigens by innate phagocytic cells (Kaushansky et al., 2013; Kurup et al., 2019; Marques-Da-Silva et al., 2021) but type I IFN can inhibit caspase activity and inflammasome activation thus potentially inhibiting the presentation of malaria antigens (Guarda et al., 2011; Veeranki et al., 2011). IFNAR signaling during malaria liver infection may also impair the induction of Th1 and CD8 T cell responses and enhance exhaustion of liver resident CD8 T cells (Haque et al., 2014; Minkah et al., 2019; Nagai et al., 2003). In addition, type I IFN can hamper protective immunity via inhibition of IFNγ responsiveness by monocytes and macrophages that in turn can limit the induction of Th1 responses (de Paus et al., 2013; Rayamajhi et al., 2010). Overall IFNγ is an important mediator in anti-malarial immunity with a variety of downstream effects besides macrophage activation (King and Lamb, 2015). We did not find that the best known and potent producers of large quantities of type I IFN, namely plasmacytoid DCs (pDCs), had significantly higher levels of in NP subjects; however, other innate cells, e.g. neutrophils, monocytes and DCs can secrete type I IFN (Ali et al., 2019; Rocha et al., 2015).

NK cell-associated transcriptome responses (Figs 3A and 4A) and relative changes in NK cell subset numbers (Fig 6B) occur earlier in P than in NP subjects, which may indicate that they contribute to the development of immunity, perhaps via the balance between Th1 and Th2 responses. NK cells can be activated by neutrophils as well as inflammatory monocytes through type I IFN (Costantini and Cassatella, 2011; Lee et al., 2017) and they can have diverse functions. They can act as immune regulators that enhance or suppress adaptive responses and they can be innate effectors that rapidly respond to and eliminate infected or tumor cells (Schuster et al., 2016; Vivier et al., 2008). IFNy secretion by NK cells can support DC-mediated Th1 induction (Schuster et al., 2016; Zhao et al., 2020), analogous to that we observed in P subjects, but NK cells can also limit adaptive responses by suppressing DCs, CD4 T cells and B cells and thus variably impact outcomes (Hayakawa et al., 2004; Piccioli et al., 2002; Zhang et al., 2013; Zhao et al., 2020). Furthermore, NK cells can reduce inflammation, e.g. as with COVID-19 related immune pathology (Li et al., 2020; Mehta et al., 2020; van Eeden et al., 2020; Zheng et al., 2020). Thus, the early NK responses in P subjects may support the priming of a Th1 polarized response and inhibit inflammation that in NP subjects primes a Th2 response. Interestingly, we identified a novel NK cell subset that expressed CD8 and lacked FcRγ expression and which is more abundant in NP subjects in the day 3-7 interval (Fig. 5D). FcRγ-lacking NKs have previously been associated with an adaptive phenotype that is protective against seasonal malaria infection (Hart et al., 2019). Further investigations into this phenotype could elucidate its functionality.

Overall, our analyses indicate that early innate responses to live irradiation attenuated SPZ vaccination substantially impact the development of adaptive responses and ultimately protection from malaria infection. Understanding mechanisms of protection is complicated by the likelihood that protective effector processes are multi-functional, due to the large breadth of potential antigens, and protection may occur at multiple points between the introduction of SPZs and the establishment of a blood stage infection. Both antibody and cellular mediated mechanisms may contribute to protection: monoclonal antibodies derived from attenuated SPZ vaccination can protect humanized mice although antibody levels variably correlate with protection (Epstein et al., 2017; Hickey et al., 2020) and liver resident CD8+ T-cells and IFNγ correlate with protection in non-human primates (Pichyangkul et al., 2017). The complex balance of responses associated with protection are illustrated here by the differential early inflammatory responses between P and NP subjects and the potential effects on Th1 and Th2 responses. The events following the priming vaccination that correlated with protection are early NK associated responses in the context of limited inflammatory responses followed by CD4+ T cell responses and subsequently CD8+ T cell responses.

That this study was performed on blood samples, despite decisive immune events occuring in the liver, which is essentially experimentally inaccessible, limited us to indirect analysis of phenotypic differences between the P and NP subjects rather than functional assays. The sample size is relatively small, which reduced our sensitivity to discover more subtle changes associated with protection, especially given natural variation in the participants. However, the detection of robust correlates of protection despite the low numbers increases our confidence that we have identified meaningful responses. These findings can inform further studies to extend the understanding of protective immunity and its development. In addition, multiple factors may have influenced the differences that between P and NP subjects that we described here, and which influenced protection. These include intrinsic differences between subjects, such as HLA type and other genetic differences as well as baseline immune status at the time of immunization. Also vaccination via infected mosquito bites may have contributed to variability in the effective vaccine dose received by each participant, i.e. the number of liver cells infected by live attenuated SPZs. In addition, the point at which parasite development in the liver cells was arrested may have been variable since radiation damage is random which may have impacted the amount and type of parasite antigen available for presentation.

In conclusion, we show that a strong acute inflammatory response to a priming vaccination correlates with the ultimate lack of protection in this trial of malaria naive volunteers. We hypothesize that this results in skewing adaptive responses toward Th2-centered responses rather than protective Th1 responses and that this similarly impacts responses to subsequent immunizations. Thus, immune responses to the first vaccination can be decisive for the outcome of the trial.

## Materials and Methods

### Sample collection and RNA sequencing

Whole blood was collected from IMRAS trial participants directly into PAXgene blood RNA tubes (PreAnalytiX, Hombrechtikon, Switzerland) and stored at -20 °C. RNA extraction and globin transcript depletion (GlobinClear, ThermoFisher Scientific, MA, USA) were performed prior to cDNA library preparation using the Illumina TruSeq Stranded mRNA sample preparation kit (Illumina, CA, USA). Globin transcript depletion, cDNA library preparation and RNA sequencing were performed by Beijing Genomics Institute (Shenzhen, China). A total of sixty-six RNA-seq samples were sequenced, with a target depth of 30 million reads per sample. Eleven of the samples were sequenced on Illumina (San Diego, CA) Hiseq2000 sequencers using 75 base-pair (bp) paired-end reads. The remaining one hundred and eighty-six samples were sequenced on BGI500 sequencers using 100 bp paired-end reads.

### Quality control and processing of RNA-Seq data

RNA-seq data were processed as previously described (Thompson et al., 2017). Read pairs were adjusted to set base calls with phred scores < 5 to ‘N’. Read pairs for which either end had fewer than 30 unambiguous base calls were removed, a method that indirectly removes pairs containing mostly adaptor sequences. Read pairs were aligned to the human genome (hg19) using STAR (v2.3.1d) (Dobin et al., 2013). Gene count tables were generated using htseq (v. 0.6.0) with the intersection-strict setting on and Ensembl gene annotations (GRCh37.74) used to link genomic locations to gene identifiers (Anders et al., 2015). Log2-transformed TMM-normalized counts-per-million (CPM) expression matrices were computed using the cpm function of the edgeR package (McCarthy et al., 2012). Batch correction for sequencer model (Hiseq2000vs BGI5000) was performed on log2-transformed counts using linear mixed-effects models with normally distributed errors and an unstructured covariance matrix. Mixed-effects models were fit using the R (https://www.r-project.org/) Lme4 package (Bates et al., 2015). The following formula was used:

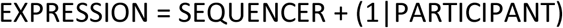

in which EXPRESSION represents the log2-transformed CPM value, SEQUENCER the sequencing platform, and including random intercepts for each PARTICIPANT. To create a final batch-corrected expression matrix, raw CPMs were adjusted by subtracting the fitted SEQUENCER coefficient.

### Mixed-effects modeling to identify transcriptional signatures that were regulated by the primary vaccine and responded differentially in protected and non-protected immunized participants

Linear mixed-effects regression models (LMER) were used to model individual gene expression (EXPRESSION) as a function of sample collection time (TIME) and protection after CHMI (PROTECTION), with TIME and EXPRESSION as fixed effects, and PARTICIPANT as a random effect.

Mixed-models were fit as follows:

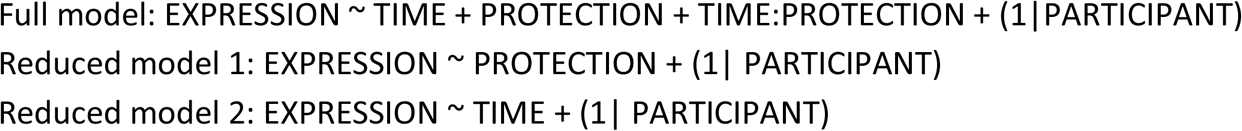

By contrasting the full model with reduced models lacking the TIME and PROTECTION terms, the significance of relationships between the TIME and PROTECTION variables and EXPRESSION were evaluated. ANOVA was used to compare the full model with reduced model 1, where *P*-values represent the significance of the improvement of fit associated with the TIME term in the analysis. FDR-adjusted *P*-values were computed using the Benjamini-Hochberg method. PROTECTION-associated genes were similarly identified within the TIME significant genes using ANOVA to compare the full model with reduced model 2.

To identify genes with significant changes in EXPRESSION at specific time points relative to the pre-vaccination state, five full models were fit for each gene with two time points (each time point following vaccination 1 and its previous time point, i.e. time intervals) included in the TIME term. In addition, for each gene, models were fit that included all time points (Days 0,1,3,7,14,28) to identify transcriptional signatures that had temporal effects at any time point.

To determine the direction (UP/DOWN) of transcriptional responses relative to either pre-vaccination time point or the previous time point in all immunized subjects, 90% confidence intervals were estimated for the TIME coefficient of the reduced model 2 as above. Cases where the lower CI > 0 were considered UP genes, upper CI < 0 were considered DOWN genes.

### Mixed-effects modeling to identify cell types that had significantly different cell proportion changes in P and NP immunized participants after the primary vaccination

Linear mixed-effects regression models (LMER) were used to model individual cell type proportion (PERCENTAGE) as a function of sample collection time (TIME) and protection after CHMI (PROTECTION), with TIME and PERCENTAGE as fixed effects, and PARTICIPANT as a random effect.

Mixed-models were fit as follows:

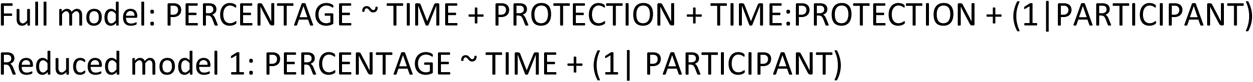

By contrasting the full model with reduced models lacking the PROTECTION terms, the significance of relationships between the PROTECTION variables and PERCENTAGE were evaluated. ANOVA was used to compare the full model with reduced model 1, where *P*-values represent the significance of the improvement of fit associated with the PROTECTION term in the analysis.

### Gene set enrichment analysis (GSEA)

GSEA was performed for each vaccination time interval using the R fgsea package (Korotkevich et al., 2016; Mootha et al., 2003; Subramanian et al., 2005) with 500 permutations and whole blood transcriptional modules (Chaussabel et al., 2008b; Li et al., 2014b; Subramanian et al., 2005; Tran et al., 2019). Genes were ranked by average fold change across each time interval (day 1 to day 0, day 3 to day 1, day 7 to day 3, day 14 to day 7, day 28 to day 14) separately for samples from P and NP subjects. Normalized enrichment scores (NES) of non-significant modules (FDR-adjusted *P*-value > 0.05) were set to 0.

### Statistical tests

The hypergeometric test was used to identify BTM modules enriched for subsets of genes. The resultant effect size (ES) was calculated as: (b/n)/(B/N), in which n: Number of genes of interest; N: Number of total mapped genes; b: Number of genes of interest from the given module; B: Number of genes from the module in the total mapped genes.

Ingenuity pathway analysis (IPA) was performed using the IPA software from Qiagen. *P*-values were calculated using Fisher’s Exact Test and FDR-adjusted P-values < 0.1 were considered significant.

### Unsupervised clustering

Hierarchical clustering of summary measures representing gene expression/responses (average gene fold changes, GSEA NES, or ES scores) was computed by agglomerative complete linkage with 1 - (Pearson’s correlation coefficient) as the distance metric. The optimal number of clusters was determined by the “elbow” method (Thorndike, 1953).

### Flow cytometry data

PBMCs were collected from IMRAS participants on day 0, and 3, 7 and 14 after the first immunization and frozen for later use. After thawing in RPMI supplemented with 10% FBS and benzonase nuclease (Millipore EMD 0.05 U/ml), the samples were incubated with LIVE/DEAD™ Fixable Blue Dead Cell Stain Kit and the Human BD Fc Block for 30 min at room temperature before being simultaneously stained with four phenotyping panels that have been previously described in OMIP-044 and OMIP-064 and further described in Table S2 (Hertoghs et al., 2020; Mair and Prlic, 2018). The cells were then acquired using a BD FACSymphony flow cytometer. The data were analyzed, and cellular populations gated and quantified using FlowJo Software (version 9.6.6). The percentage contribution of each manually gated cell subset was calculated using the counts of each defined cell subset divided by the total single live cells from that sample. Pearson correlation coefficients were calculated between cell type proportion changes and BTM mean expression level changes per-time interval for P an NP subjects separately.

### CITE-seq single-cell RNA seq processing

Live frozen PBMCs were obtained from a single vaccinated individual in cohort 1 of the IMRAS trial at day 0, and three- and 14-days post first vaccination. Cells were thawed and washed with RPMI supplemented with 10% FBS and benzonase nuclease (Millipore EMD 0.05 U/ml). PBMCs were resuspended in 100 µl of PBS supplemented with 2% w/v Fetal Bovine Serum (FBS) and incubated with Fixable Viability Stain 510 and Human BD Fc Block for 30 minutes at room temperature. Cells were washed with 2% FBS PBS before incubating with a panel of previously titrated 14 barcoded oligo-conjugated antibodies (BioLegend TotalSeq-C), including FITC-anti-CD45. Stained PBMC samples were then sorted by fluorescence activated cell sorting (FACS) on a BD FACSMelody to enrich for live, hematopoietic cells. A standard viable CD45+ cell gating scheme was employed; FSC-A v SSCA (to exclude sub-cellular debris), two FSC-A doublet exclusion gates (FSC-W followed by FSC-H), dead cell exclusion gate (BV510 LIVE/DEAD negative) followed by CD45+ inclusion gate.

Sorted cells were resuspended in PBS supplemented with 1% BSA. Cells were loaded onto the 10X Chromium system, where we aimed for recovery of ∼5000 cells per sample, and subjected to partitioning with barcoded 5’ V1.1 chemistry gel-beads (10X Genomics) to generate the Gel-Bead in Emulsions (GEMs). The RT reaction was conducted in the GEMs, barcoded cDNA extracted by post-GEM RT-cleanup, and cDNA and antibody barcodes amplified with 14 cycles. Amplified cDNA was subjected to SPRI bead cleanup at 0.6X. Amplified antibody barcodes were recovered from the supernatant and were processed to generate TotalSeq-C libraries as instructed by the manufacturers (10X Genomics and BioLegend, TotalSeq-C with 10x Feature Barcoding Protocol). The remaining amplified cDNA was subjected to enzymatic fragmentation, end-repair, A-tailing, adapter ligation and 10X specific sample indexing as per manufacturer’s protocol. Libraries were quantified using Bioanalyzer (Agilent) analysis. 10x Genomics scRNA-Seq and TotalSeq-C libraries were pooled and sequenced on an Illumina NovaSeq Sp100 flow cell using the recommended sequencing read lengths of 26 bp (Read 1), 8 bp (i7 Index Read), and 91 bp (Read 2), and depths of 50,000 and 5000 read pairs per cell for the 5’ Gene Expression and TotalSeq-C libraries respectively. Cell Ranger v3.1.0 (10x Genomics) was used to demultiplex raw sequencing data and quantitate transcript levels against the 10x Genomics GRCh38 reference.

### Single-cell RNA seq processing and analysis

Raw count data were filtered to remove cells where 1) a mitochondrial RNA fraction greater than 7.5% of total RNA counts per cell, and 2) less than 200 or greater than 2500 genes were detected. The resultant count matrix was used to create a Seurat (v4.0.1) (Hao et al., 2021) object. Filtered read counts were normalized, scaled, and corrected for mitochondrial and rRNA read percentages with the SCTransform function. The ADT matrix was normalized per feature using center log normalization. Cell types in each sample were annotated by mapping to the annotated reference PBMC dataset provided in the Seurat v4 Azimuth workflow. Briefly, anchors between the query and reference datasets were identified using a precomputed supervised PCA on the reference dataset. Next, cell type labels from the reference dataset, as well as imputations of all measured protein markers, were transferred to each cell of the query datasets through the previously identified anchors. The query datasets were then merged and projected onto the UMAP structure of the reference. The genes expressed in each specific cell cluster were identified using the FindAllMarkers function from the Seurat4 package and filtered to include those with average log_2_ fold changes greater than 1 and FDR-adjusted P-values less than 0.05.

## Acknowledgements

We would like to thank Michael Gale and Nana Minkah for their helpful comments on the manuscript. We would also like to acknowledge the contribution of all the IMRAS study volunteers.

## Supplementary Material

**Figure S1. Overview of the IMRAS trial. A**. Schematic indicating timing of vaccination and sampling. **B**. Kaplan-Meier curve showing the percentage of true immunized subjects who did not develop parasitemia after CHMI (green) and those who did (red). **C**. Number of infectious mosquito bites received by each subject at each immunization. Circles and triangles indicate non-protected and protected subjects, respectively. Red lines indicate the median number of infectious mosquito bites.

**Figure S2. Comparison of protection associated clusters P_1 and NP_1. A**,**B**. Gene overlap of IPA pathways and BTMs enriched in cluster P_1 (**A**) and NP_1 (**B**), Node sizes indicate numbers of genes in a BTM or IPA pathway, and line thickness indicates the numbers of shared genes between two nodes. **C**. Heatmap showing expression of 317 genes common in cluster 1 of P subjects and cluster 1 of NP subjects. Expression values were z-score transformed in rows for visualization.

**Figure S3. Expression changes in pattern-recognition receptor pathwaysxs. A**,**B** Heat maps of expression changes for MyD88-dependent toll-like receptor associated genes (**A**) and MyD88-independent toll-like receptor associated genes (B) in P and NP subjects. Genes shown were selected using Gene-ontology (GO) annotations GO:0002755 (MyD88 dependent TLR signalling pathway) and GO:0002756 (MyD88 independent TLR signalling pathway). Expression values were Z-score transformed in rows for visualization. **C**. Average gene expression of selected IPA pathways over time in P (green) and NP (red) subjects. Dots represent average gene expression values per-individuals and solid line represents the average gene expression of the IPA pathway across all participants.

**Figure S4. Expanded color legend showing IPA pathways and BTMs enriched in Fig. 2**

**Figure S5. Expression changes in reactive-oxygen species pathway genes. A**. Heatmap showing expression of Hallmark Reactive Oxygen Species (ROS) Pathway genes. Expression values were z-score transformed in rows for visualization. **B**. STRING-DB derived protein-protein interaction networks of ROS genes, colored by expression changes on day 1 compared to day 0 seperately for P and NP. **C**. Average expression profiles of genes of Hallmark Reactive Oxygen Species Pathway in P and NP subjects. Dots represent average expression in individuals and solid line represents the average expression of the pathway over all P or NP subjects.

**Figure S6. Expression changes in pathways associated with specific immune cell types: A-I**.Heatmaps showing expression of genes of cell-type specific pathways: Neutrophils (**A**), Monocytes (**B**), DCs (**C**), Platelets (**D**), T-cells (**E**), B cells (**F**), NK cells (**G**), classically activated macrophages (**H**) and alternatively activated macrophages (**I**). Expression values were z-score transformed in rows for visualization.

**Table S1**. Genes and pathways associated with hierarchical clustering of protection-associated genes

**Table S2**. List of antibodies used for high parameter flow cytometry. Three separate staining panels for the detection of DC subsets, monocytes, T cells and NK cells are shown.

**Table S3:** List of CITEseq antibodies

