## Supplementary figures and images for "Systems analysis of immune responses to attenuated *P. falciparum* malaria sporozoite vaccination reveals excessive inflammatory signatures correlating with impaired immunity"

### Figure S1, Figure S2, Figure S3, Figure S4, Figure S5, Figure S6

A

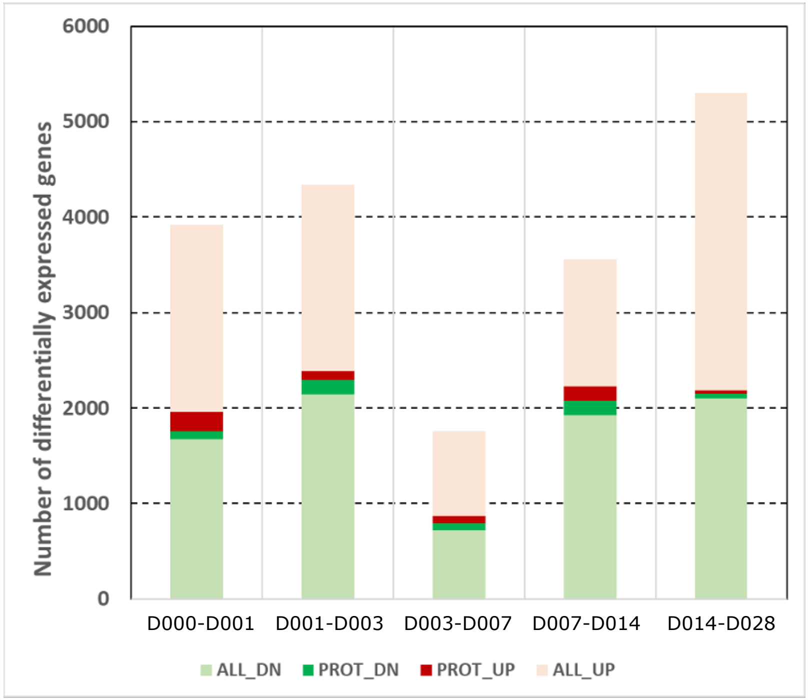


| INTERVAL | ALL_UP | ALL_DN | PROT_UP | PROT_DN |
|---|---|---|---|---|
| D000-D001 | 1961 | 1669 | 208 | 84 |
| D001-D003 | 1956 | 2138 | 91 | 152 |
| D003-D007 | 885 | 714 | 76 | 81 |
| D007-D014 | 1330 | 1925 | 148 | 151 |
| D014-D028 | 3113 | 2103 | 36 | 45 |
| #of unique genes | 8170 | | 806 | |

B

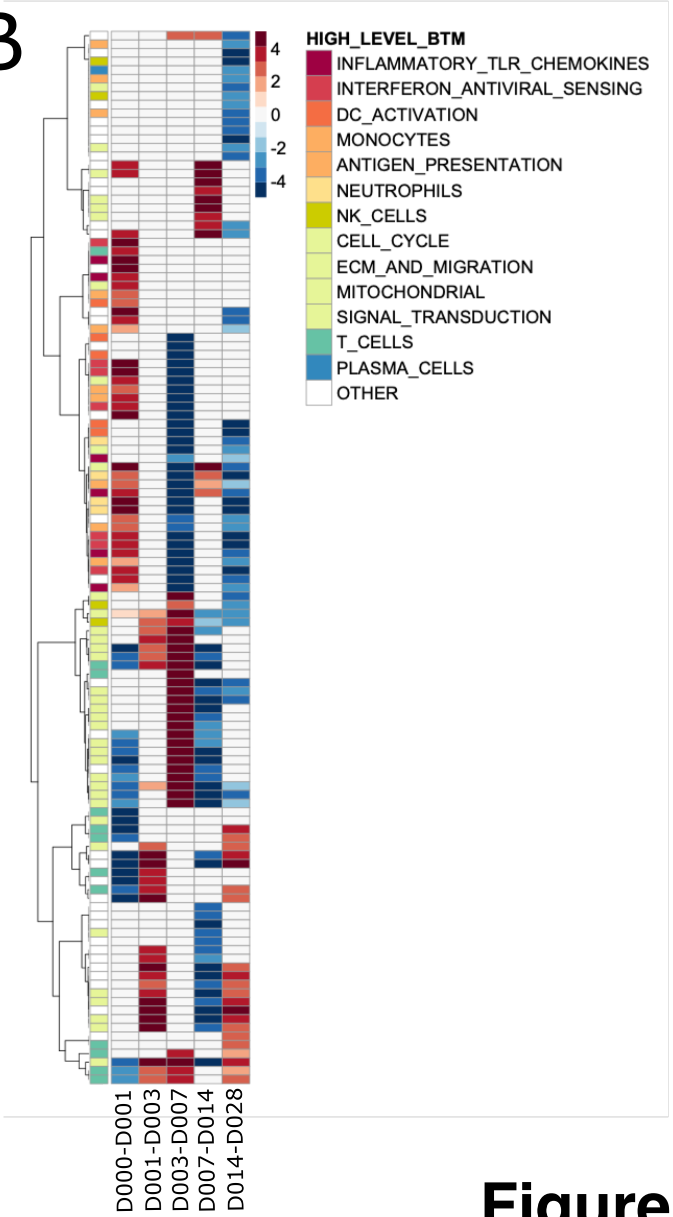


**Figure 1**

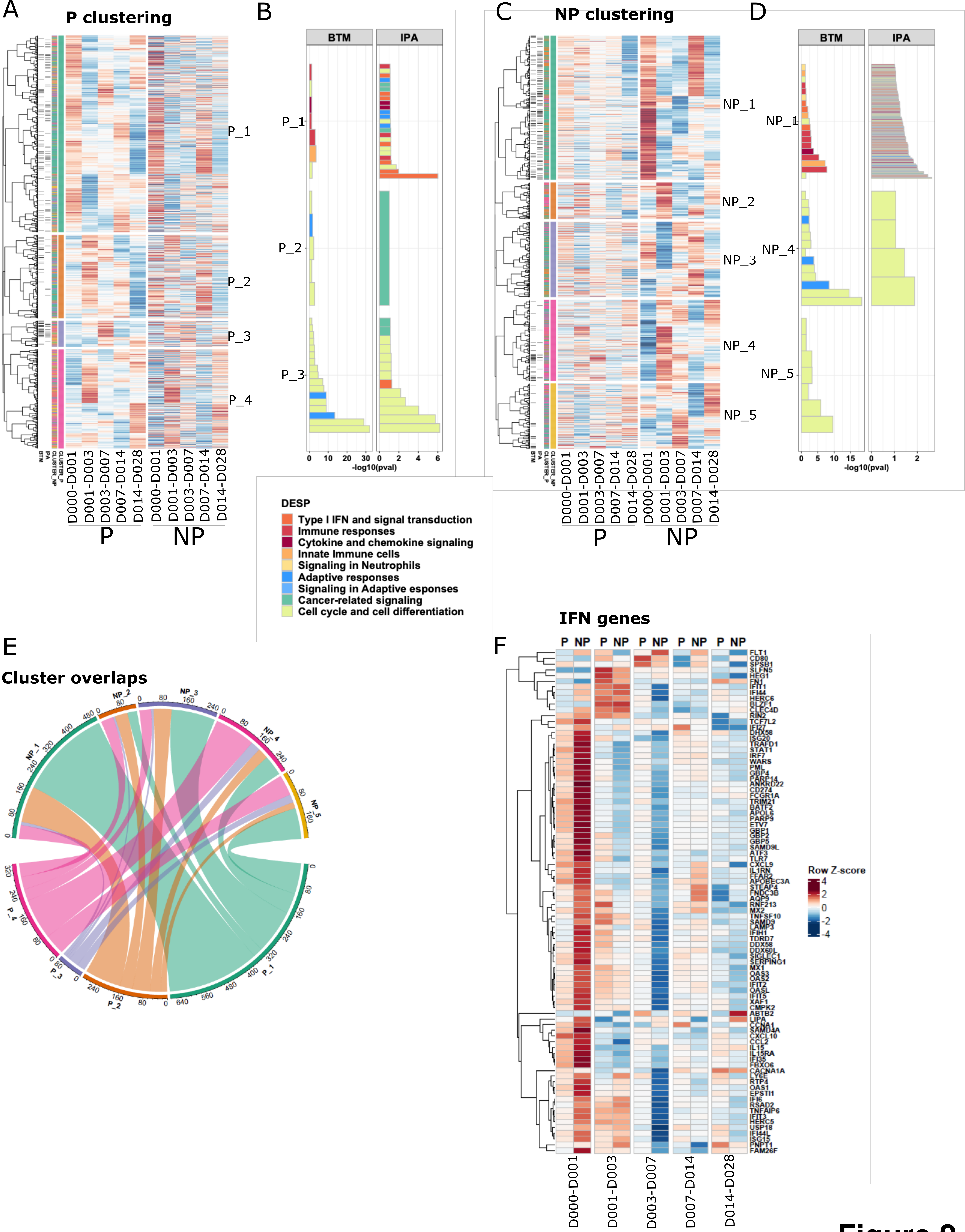

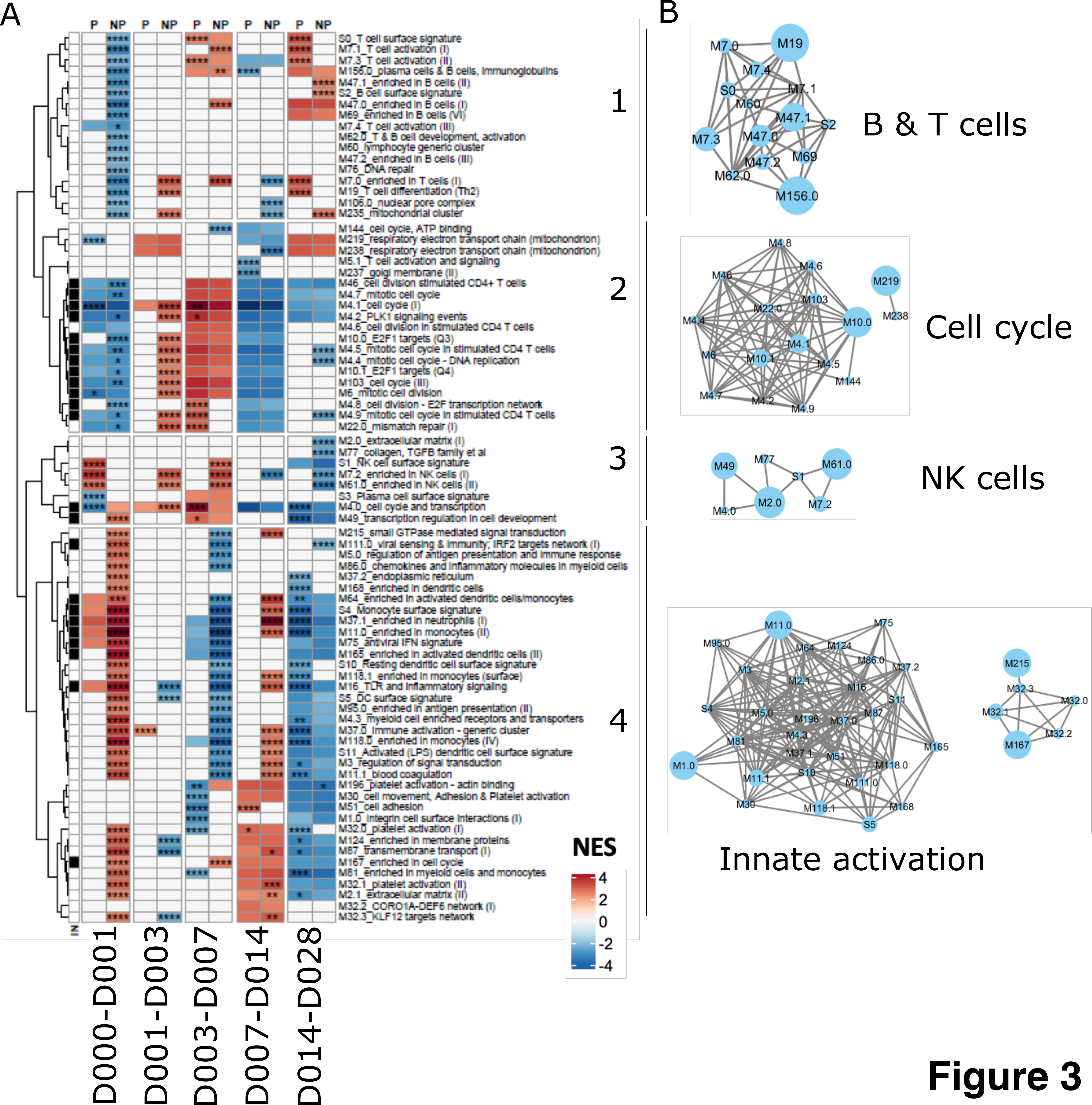

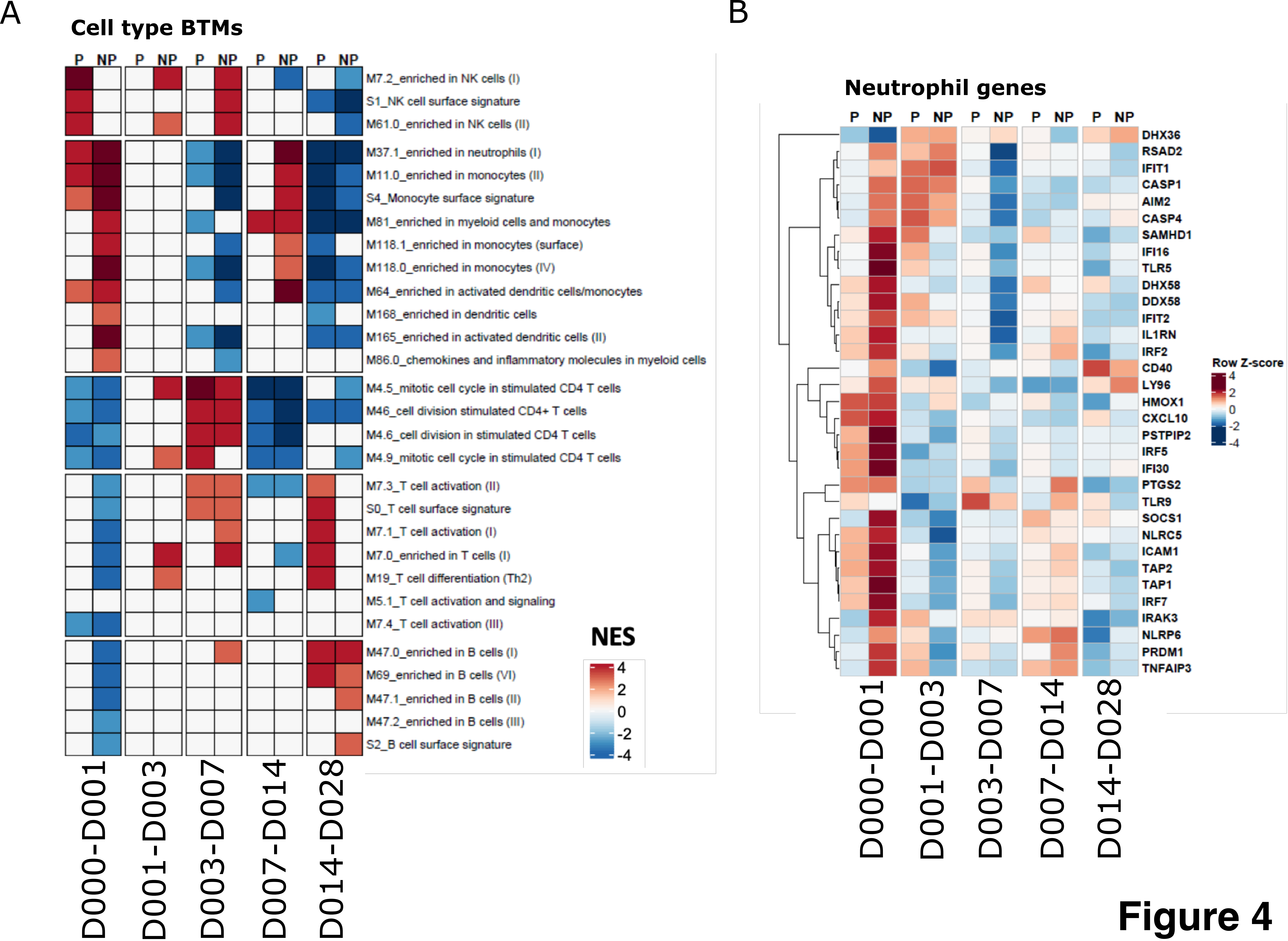

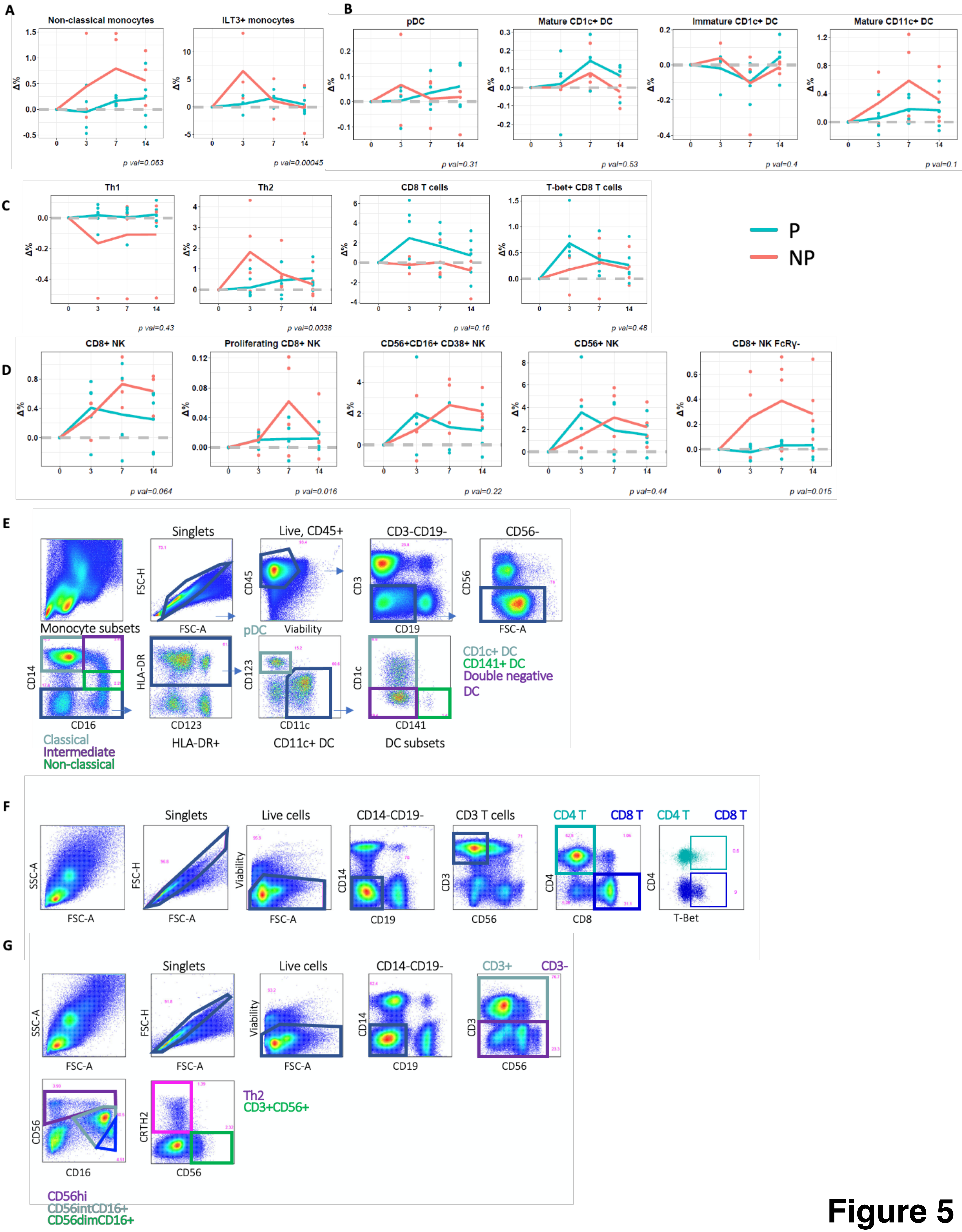

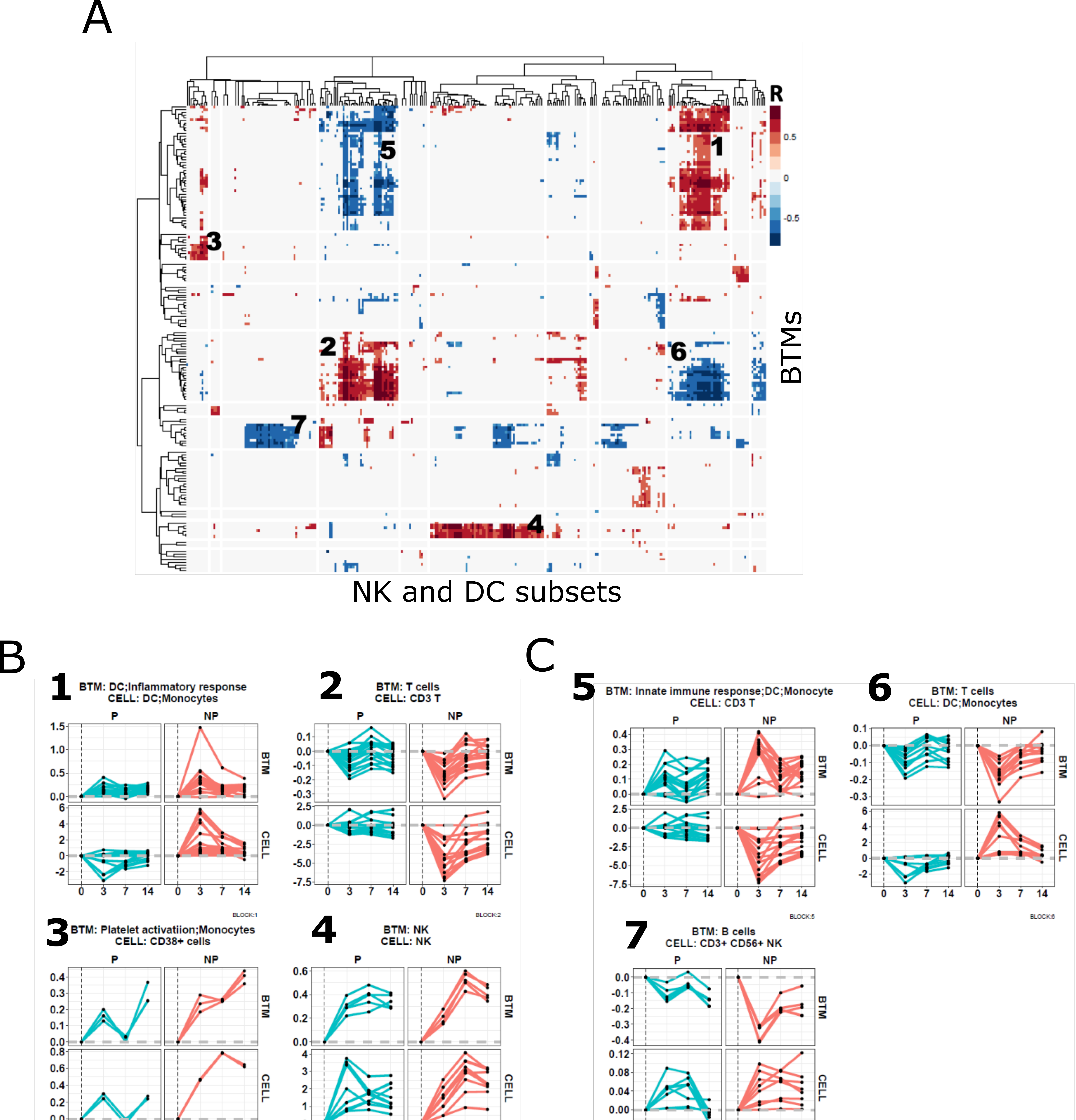

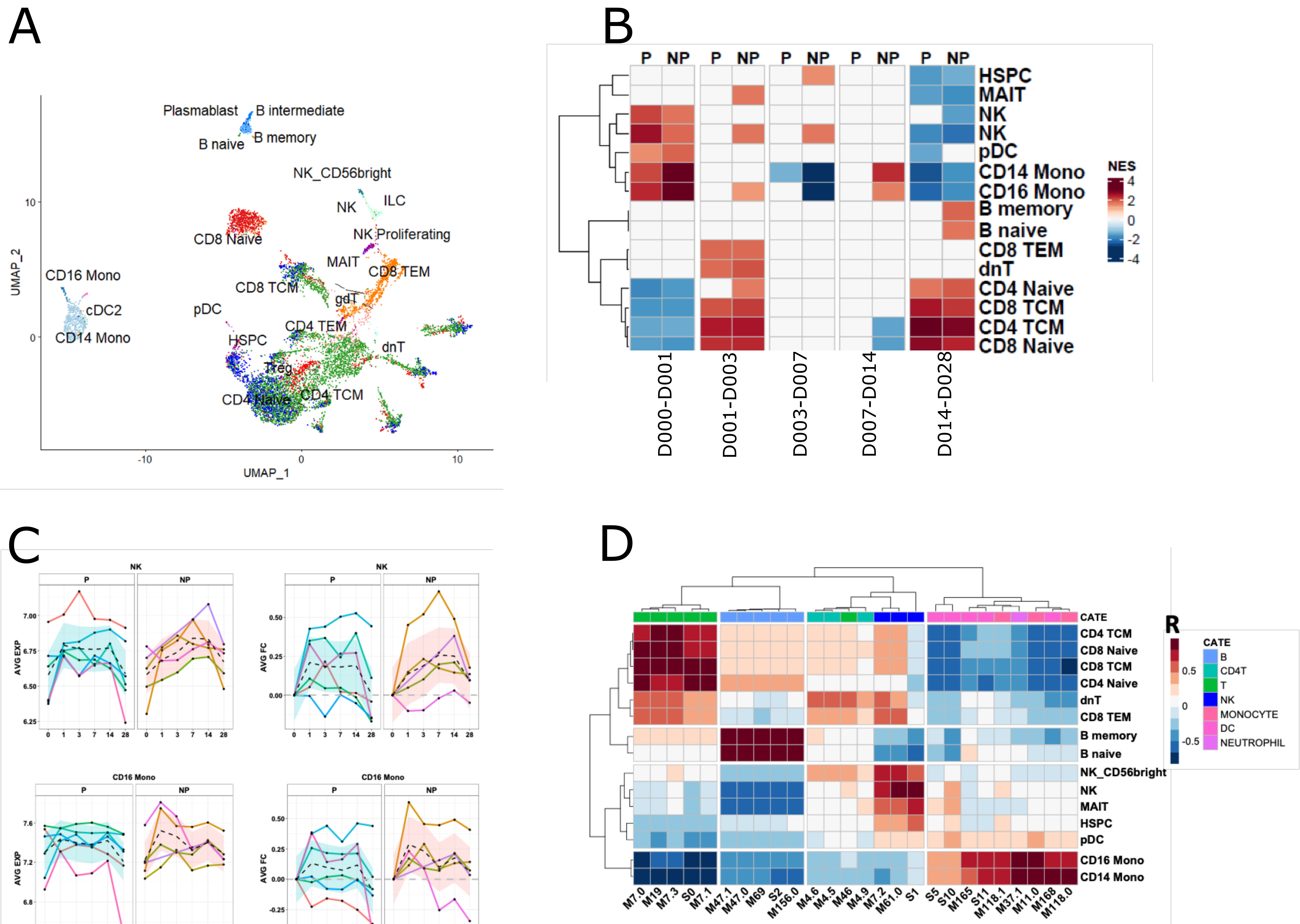
